# Exhaustive Isotope Tracing Reveals the Butterfly Effect of Mammalian Producer Cell Line Selection for Biomanufacturing

**DOI:** 10.64898/2026.08.04.742784

**Authors:** Venkata Gayatri Dhara, Jacqueline E. Gonzalez, Harnish Mukesh Naik, Brian O. McConnell, Pratik Anay Khare, Justin Sargunas, Maciek R. Antoniewicz, Michael J. Betenbaugh

## Abstract

Chinese hamster ovary (CHO) cells are the dominant platform for recombinant biotherapeutics, yet the impact of producer cell line selection on mammalian cell metabolism remains poorly understood. Here, we performed 41 independent ^13^C-tracer experiments using uniformly labeled glucose or individual amino acids to comprehensively map carbon utilization in the two principal CHO production platforms: methotrexate-selected CHO-K1 and glutamine synthetase-selected CHO-GS cells. Time-resolved GC-MS analysis revealed distinct metabolic phenotypes spanning central carbon metabolism, amino acid interconversion, lipid biosynthesis, and one-carbon metabolism. CHO-K1 cells exhibited extensive reductive carboxylation and pyruvate carboxylase-mediated anaplerosis, whereas CHO-GS cells redirected glutamate toward glutamine synthesis and relied on asparagine and aspartate to support TCA cycle activity. Isotopomer analysis uncovered substantial intracellular-extracellular cycling of alanine, glycine, glutamate, and serine despite contrasting uptake profiles and quantified differential amino acid contributions to fatty acids and cholesterol. Serine, glycine, and methionine labeling revealed active folate-cycle interconversion in CHO-K1 and enhanced methionine-cycle activity in CHO-GS. Aspartate is identified as a key redox exchange factor and uniquely informative tracer for pathway characterization following glutamine depletion. Together, this exhaustive isotope-tracing framework establishes how producer cell line selection rewires mammalian metabolism and provides a foundation for cell engineering, media optimization, metabolic modeling, and next-generation biomanufacturing.

## 1. Introduction

The proportion of biotherapeutics in the global pharmaceutical market has expanded steadily with more than 400 biologic drugs approved since 2002, including monoclonal antibodies (mAbs), vaccines, fusion proteins, and bispecific antibodies (BsAbs) ^1,2^. Mammalian cell lines are the principal drivers for biologics manufacturing ^3^ with Chinese hamster ovary (CHO) cells responsible for the bioproduction of nearly 70% of all approved biologics ^2,4^. CHO cells produce recombinant proteins with high productivities, human-compatible glycosylation patterns, and exhibit scalability into large suspension bioreactor formats ^5^. However, the growing demand globally for biologics from CHO and their comparatively high bioproduction costs drives innovation to increase biomanufacturing efficiencies and lower bioproduction costs through advances in cell line engineering, enhanced process analytics and control, and optimization of cell culture media and feed formulations.

A critical factor in designing media and feeds for biomanufacturing platforms is knowledge of the utilization of specific media components, in particular amino acids and glucose, carbon sources that represent the lion’s share of nutrients consumed, as well as the metabolic pathways these nutrients follow to generate biomass components. This is especially relevant in CHO and other mammalian cells because of their utilization of different sources of carbon and nitrogen, which can follow multiple metabolic pathways, ultimately reaching fates that contribute to energy generation for growth and production of metabolic components of biomass and biologics. Understanding how CHO cells uptake and utilize amino acids and glucose is therefore crucial to develop basal media and feeds needed for optimal cell growth and efficient biomanufacturing processes.

Producing these high value biotherapeutics, including monoclonal antibodies and other recombinant proteins, requires the stable integration of one or more copies of target gene into the genome. The process typically includes transfection of the gene of interest together with a selectable marker, followed by selection, gene amplification (in some cases), and clone screening to isolate cell lines producing the gene of interest. The two most widely used CHO selection platforms employ either the dihydrofolate reductase (DHFR) or glutamine synthetase (GS) selection systems. In the DHFR system, gene amplification is driven by increasing concentrations of methotrexate, resulting in multiple copies of both the DHFR gene and the linked transgene ^6,7^. In contrast, the GS system relies on glutamine deprivation and inhibition of endogenous glutamine synthetase activity, often using methionine sulfoximine (MSX), to enrich for cells expression levels ^8,9^.

Developing optimal media formulations and biomanufacturing conditions for these recombinant CHO cell lines represents a significant challenge because each cell line may possess unique physiological and metabolic characteristics. These intrinsic differences arise not only from the parental CHO host but are often amplified by the cell line development process itself.

The very action of selective pressures imposed during DHFR- or GS-based cell line generation has the potential to alter the intrinsic cellular physiology and consequently the metabolic requirements for specific nutrients including particular amino acids. As a result, the choice of selection system and cell line development strategy has lasting effects on metabolic utilization and distributions patterns that can have far reaching impacts on processing condition and biomanufacturing performance in different environments.

Stable isotopes – including ^13^C-labeled amino acids and glucose – have become a valuable tool to unravel the fate of critical substrates in mammalian cell culture. Isotopic labels are incorporated into biomass components and various intracellular or secreted metabolites that can then be followed through analytical techniques such as GC-MS, LC-MS, and NMR ^10–13^. Only a limited number of isotopic tracer substrates have been used for labeling studies in CHO cell cultures; including glucose, glutamine, asparagine, pyruvate, and some fatty acids ^14–19^. Thus far, ^13^C-glucose and ^13^C-glutamine tracers have been most widely used for stable-isotope tracing and ^13^C-metabolic flux analysis (^13^C-MFA). Yet these tracers provide only one window into metabolism; there is a dearth of data on other cellular substrates, including many of the 20 amino acids in typical culture medium. This is surprising given the importance of balancing these components to create formulations supporting robust growth in CHO and other mammalian bioproduction platforms. Isotope labeling can play a unique, useful role in understanding the metabolic disposition of substrates and the changes in metabolite pools as the cells progress through growth phases, revealing shifts in the cell state over the course of cell culture processes. This knowledge can be invaluable in the design of new and improved media formulations, validation of cellular metabolic models, and optimization of cellular performance for maximizing growth and recombinant protein production.

In this work, we applied ^13^C-labeling of glucose and 20 amino acids to establish the metabolic architecture of CHO cells. Specifically, we evaluated and compared metabolic phenotypes between these two most widely used and prototypical CHO cells lines used in biomanufacturing: CHO-K1 and CHO-GS. To accomplish this goal, we generated complete cell culture medium deficient only in glucose or a single specific amino acid, then supplemented a fully ^13^C-labeled version of that missing component (for example, fully labeled ^13^C-leucine was supplemented to leucine-deficient cell culture medium) (**Fig 1**). From these exhaustive tracking studies, we were able to determine the fate and culture dynamics of the vast majority of amino acids in CHO cell metabolism as well as determine how the cell distributed its available resources to metabolites, lipids, and biomass components. Mass isotopomer distributions for identified key metabolites yielded valuable insights into how key metabolic pathways – glycolysis, the citric acid cycle, aspartate-malate shuttle, lipogenesis, one carbon metabolism, and so forth – differed between the two principal CHO cell platforms. This study thus represents by far the most exhaustive analysis of amino acid utilization of mammalian bioproduction system to date and provides a transformative understanding of how the cell line selection itself defines the metabolic architecture of producer cells with consequential impacts on utilization of key nutrients, production rates of key metabolites and ultimately the ability to develop an optimized biomanufacturing process for the next generation of biologics emerging from this dominant bioproduction workhorse of the biotechnology industry.

**Figure 1.**
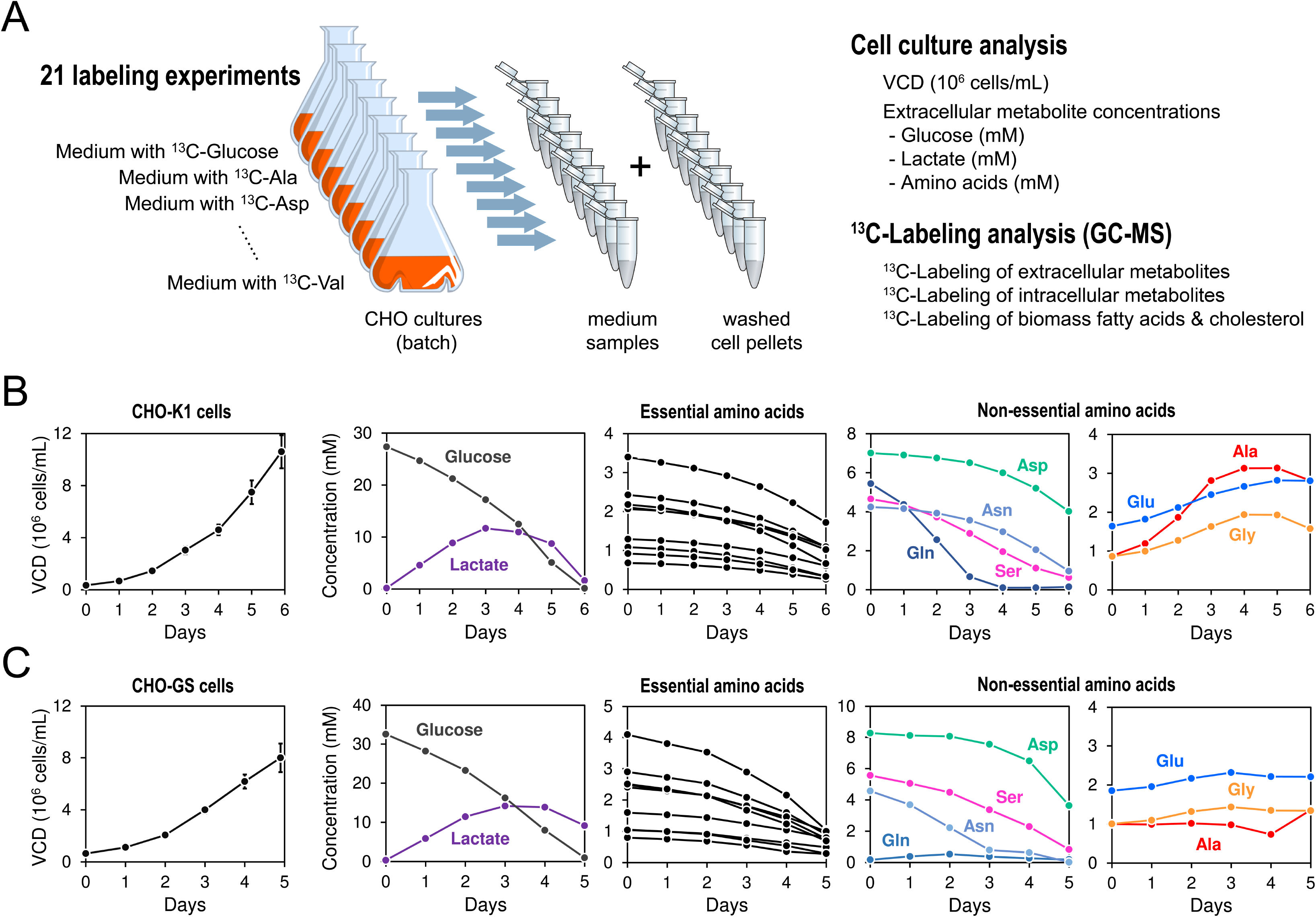
Stable-isotope labeling experiments to trace the metabolism of glucose and amino acids in two CHO cell lines. (A) CHO cells were grown in 21 parallel cultures using custom media, each containing a different ^13^C-tracer, either glucose or an amino acid. On each day of the culture, cell growth and the uptake and secretion of extracellular metabolites were measured. ^13^C-Labeling incorporation was also measured daily by GC-MS for extracellular metabolites, intracellular metabolites, fatty acids and cholesterol. (B) Viable cell density (VCD) and extracellular metabolite concentrations for CHO-K1 cell cultures, and (C) CHO-GS cell cultures. All measured data are provided in supplemental materials.

## 2. Results

### 2.1. Physiological characterization of CHO-K1 and CHO-GS cell cultures

A total of 41 comprehensive ^13^C-tracer experiments were conducted across the two previously established CHO cell lines– CHO-K1 and CHO-GS – to trace glucose and amino acid metabolism and uncover phenotypic differences between these two prototypical platforms. In each tracer experiment, a different substrate in the medium was replaced with a fully ^13^C-labeled version of that component (**Fig. 1A**). For the CHO-GS cell line, ^13^C-glutamine studies were not conducted due to the integration of the Glutamine Synthetase (*GS*) gene, which allows for the synthesis of glutamine from glutamate and ammonia, allowing the cells to be cultured in glutamine-free medium with no extracellular glutamine available for replacement with fully ^13^C-labeled glutamine.

To characterize the physiology of these CHO cells in culture, the viable cell density (VCD) and concentrations of glucose, lactate, and amino acids were monitored each day over a batch culture (**Fig. 1B, C**). Both cell lines exhibited exponential growth in early culture to peak viable cell densities of 10.6 and 8 x 10^6^ cells/mL for CHO-K1 and CHO-GS, respectively, with glucose nearly fully consumed by day 6 for CHO-K1 and day 5 for CHO-GS. Lactate was produced in early culture, then gradually consumed starting on day 3 for CHO-K1 and day 4 for CHO-GS, exhibiting a lactate shift observed in many CHO cell cultures ^20–22^. For both CHO cell lines, essential amino acids were consumed at rates that met the requirements for cell growth, including proline, for which both CHO cell lines used in this study are auxotrophic ^23^. Three non-essential amino acids (asparagine, aspartate, and serine), as well as glutamine for CHO-K1 cells, were consumed at significant rates, suggesting these amino acids were likely catabolized by CHO cells or interconverted into other metabolites. Glutamine was nearly fully consumed by day 3 by CHO-K1 cells, while asparagine was effectively exhausted by day 3 by CHO-GS cells. The concentrations of glutamate, glycine, and alanine increased for CHO-K1 cells (**Fig. 1B**), while they remained relatively stable for CHO-GS cells (**Fig. 1C**), clear indicators of metabolic differences between cell lines. Concentration profiles for these amino acids and metabolites have been tabulated in **Supplemental Tables 2 and 3**.

### 2.2. Tracing the dynamics of carbon flow from amino acids and glucose through extracellular and intracellular central carbon metabolite pools

The percentage of ^13^C-labeling in extracellular and intracellular metabolites was determined for all 41 tracer experiments using gas chromatography mass spectrometry (GC/MS). This allowed for the dynamic tracking of extracellular amino acids and three extracellular organic acids (lactate, pyruvate, and citrate) for the two CHO cell lines (**Fig. 2**). Each column of **Fig. 2** represents a daily ^13^C-tracer experimental measurement for a specific labeled amino acid or glucose tracer, with the rows denoting different measurable entities with significant labeling from the source tracer. A percentage labeling of higher than 2% was considered significant, and thus measurable entities were considered labeled if the percentage labeling was between 2% and 100%.

**Figure 2.**
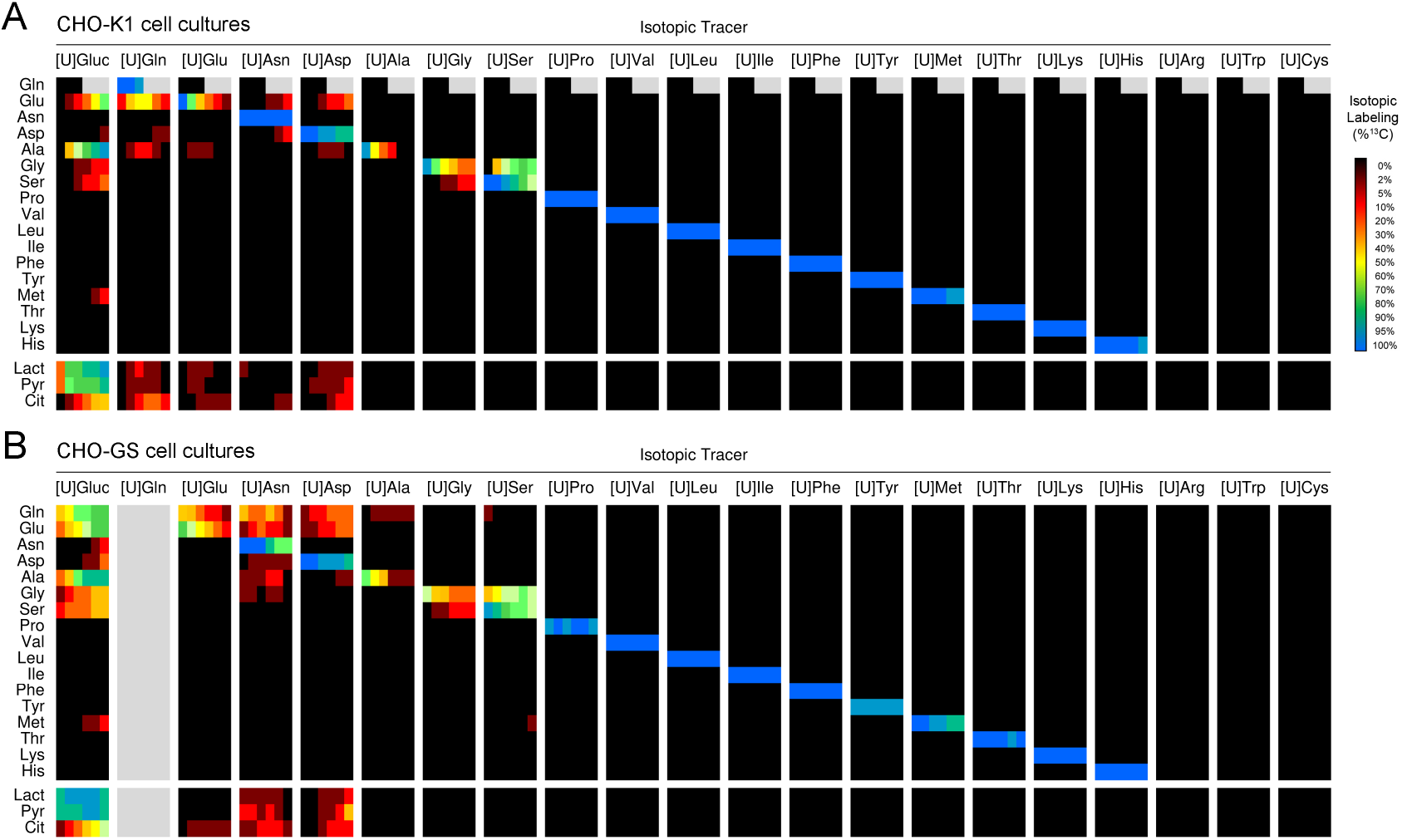
^13^C-Labeling of extracellular metabolites from parallel labeling experiments for (A) CHO-K1 cells, and (B) CHO-GS cells. ^13^C-Labeling was measured on each day using GC-MS for measurable extracellular amino acids, lactate, pyruvate and citrate. Because the medium for CHO-GS cells does not contain glutamine, only 20 labeling experiments were conducted with CHO-GS cells. To calculate isotopic labeling, the measured mass isotopomer distributions were first corrected for natural isotope abundances, and then the ^13^C-labeling was calculated as follows: isotopic labeling (%^13^C) = Σ i×M_i_. All measured data are provided in supplemental materials.

The enrichment of most secreted essential amino acids that could be detected in the medium (Pro, Val, Leu, Ile, Phe, Tyr, Thr, Lys, His) did not change significantly during the tracer experiments; in other words, the enrichment remained constant at about 100% labeling across all days (**Fig. 2**). This was expected given that essential amino acids cannot be synthesized by CHO cells from other sources. The exceptions to this trend were tyrosine, proline, and threonine in CHO-GS (**Fig. 2B**) and methionine in both cell lines (**Fig. 2A, B**), which exhibited high but not complete enrichment over the course of cell culture. Three amino acids – arginine, tryptophan, and cysteine – were not detected by our GC-MS method and could therefore not be assessed.

Glucose exhibited a significant dynamic labeling distribution into many extracellular metabolites, as did four amino acids across the two cell lines: glutamate, glutamine (in CHO-K1), asparagine and aspartate. These components contributed to varying degrees towards enrichment of a few amino acids (notably alanine), as well as the organic acids lactate, pyruvate and citrate. In both cell lines, serine and glycine contributed to each other’s extracellular labeling, with serine playing a much more prominent role in the accumulation of both amino acids (**Fig. 2**).

Similarly, the profiles of ^13^C-labeling of intracellular amino acids and metabolic intermediates in central carbon metabolic pathways were quantified (**Fig. 3**). These included glycolytic metabolites – 3-phosphoglyceric acid (3PG), phosphoenolpyruvate (PEP), pyruvate, and lactate – and citric acid cycle metabolites: citrate, succinate, fumarate, and malate. In addition to these intracellular species, we quantified ^13^C-labeling incorporation into cellular fatty acids (C16:0 and C18:0) and cholesterol, which are produced from cytosolic acetyl-CoA by CHO cells.

**Figure 3.**
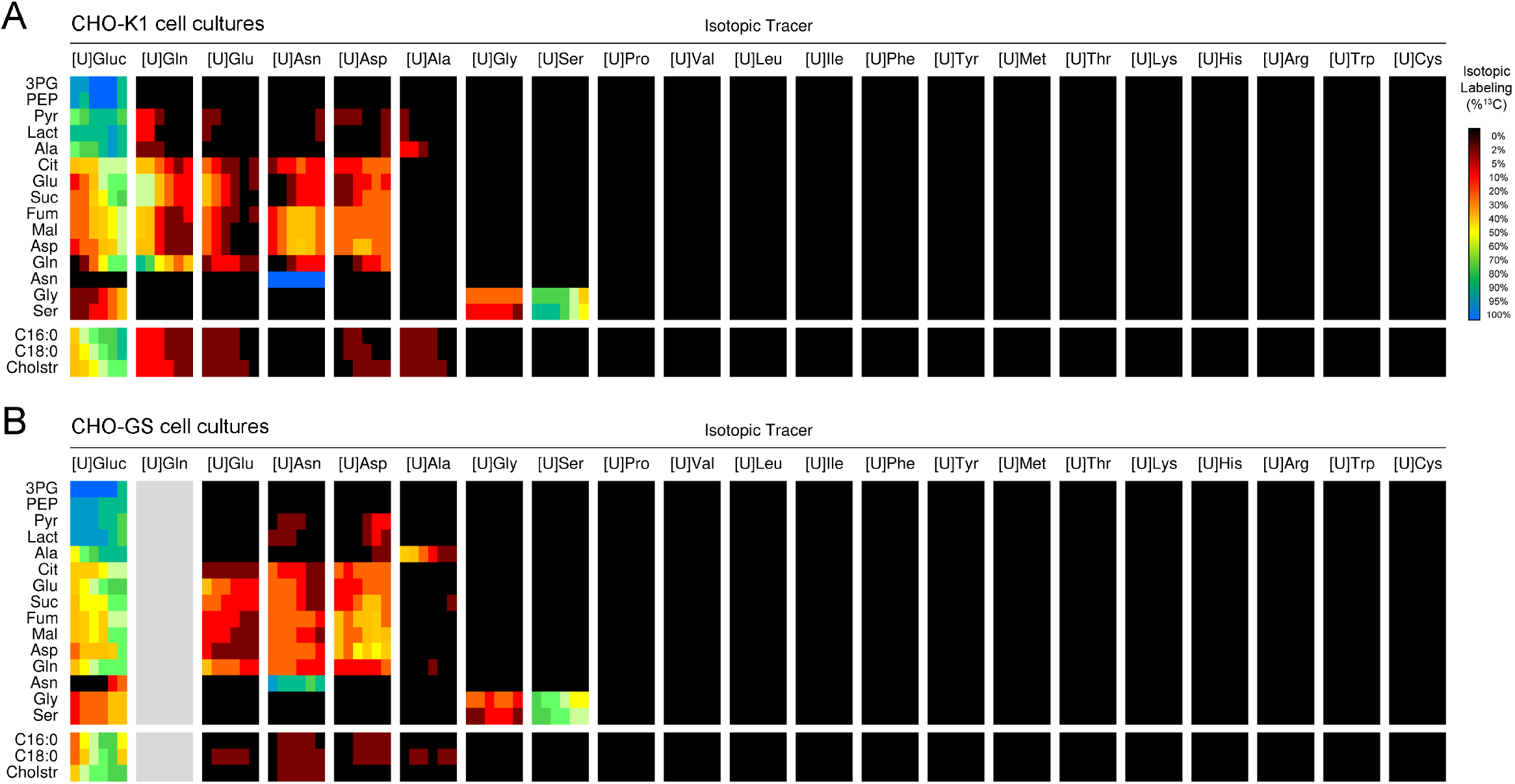
^13^C-Labeling of intracellular metabolites from parallel labeling experiments for (A) CHO-K1 cells, and (B) CHO-GS cells. ^13^C-Labeling was measured on each day for metabolites in central carbon metabolism, fatty acids, and cholesterol. Because the medium for CHO-GS cells does not contain glutamine, only 20 labeling experiments were conducted with CHO-GS cells. To calculate isotopic labeling, the measured mass isotopomer distributions were first corrected for natural isotope abundances, and then the ^13^C-labeling was calculated as follows: isotopic labeling (%^13^C) = Σ i×M_i_. All measured labeling data, including for intracellular amino acids (not shown here), are provided in supplemental materials.

Glucose was the principal contributor to many glycolytic intermediates and TCA cycle metabolites, with the same key amino acids – glutamine (in CHO-K1), glutamate, asparagine, and aspartate – also making significant contributions. We did not observe any of the essential amino acids contributing ^13^C-labeling to any of the other amino acids nor central carbon metabolites. We further identified cell-specific differences in the intracellular labeling patterns derived from several amino acids. Glutamine, which is a key metabolic nutrient in CHO-K1 cells, contributed to pyruvate, lactate, and alanine (**Fig. 3A**). While glutamate was also used minimally to form pyruvate and lactate in early CHO-K1 culture, it did not yield detectable levels of those products in CHO-GS. In CHO-GS cells, asparagine, followed by aspartate, contributed to these glycolytic metabolites along with glucose. Differences were also observed between the two cell lines in the sources of cellular fatty acids (C16:0 and C18:0). In CHO-K1, glutamine was again prominent, contributing appreciably to fatty acid labeling. Glutamate, aspartate, and alanine, but not asparagine, contributed to fatty acid synthesis in lesser amounts (**Fig. 3A**). In CHO-GS, asparagine contributed to C16:0, C18:0, and cholesterol pools; glutamate, aspartate, and alanine also exhibited limited incorporation into some of these species (**Fig. 3B**).

In CHO-GS cells, glucose contributed to asparagine synthesis during the later stages of culture, a phenomenon not observed in CHO-K1 cells. Alanine contributed modestly to lactate and pyruvate early in cultures for CHO-K1 but not in CHO-GS and was utilized differentially to synthesize fatty acids and cholesterol. Differences were also observed in alanine uptake between the two cell lines as well (**Fig. 1**), marking it as an amino acid of interest when characterizing metabolic differences between the cell lines. As with the extracellular metabolites, serine was the largest contributor to both intracellular serine and glycine in both cell lines, with minor temporal differences observed in the enrichment patterns of these amino acids.

Isotopic labeling of intracellular metabolites did not remain constant over the culture duration for both CHO cell lines. To better visualize this, we calculated the relative contributions of key carbon sources (**Fig. 4A**) to each intracellular metabolite based on the amount of labeling incorporated from the respective tracer (**Fig. 4B** for CHO-K1, **Fig. 4C** for CHO-GS). Broadly, we observed that the relative contribution of glucose steadily increased over time, as evidenced by increased labeling in intracellular metabolites from ^13^C-labeled glucose tracer (in gray), while the combined contributions of glutamine, glutamate, asparagine, and aspartate declined in the batch cell cultures.

**Figure 4.**
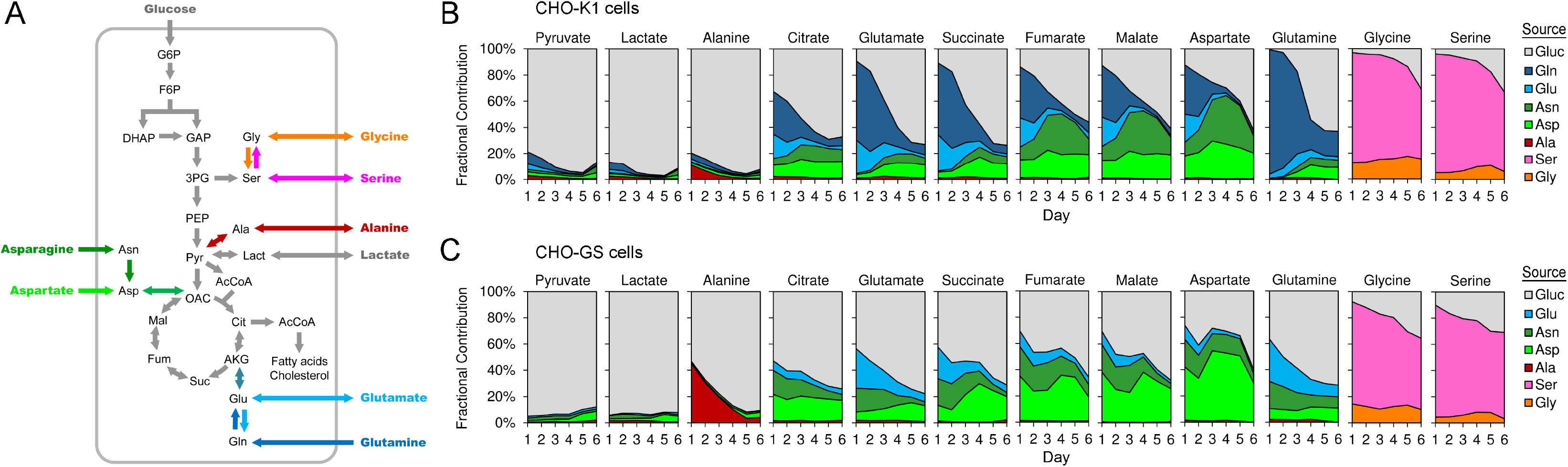
Tracing the dynamics of carbon flow from amino acids and glucose in CHO cells. (A) Diagram of biochemical pathways engaged in metabolism of glucose and amino acids. The fractional contribution of different carbon sources to the production of specific intracellular metabolites was calculated from the measured isotopic labeling in parallel labeling experiments for (A) CHO-K1 cells, and (B) CHO-GS cells.

Glucose played the dominant role in the enrichment of the glycolytic metabolites pyruvate and lactate in both cell lines, as expected. Unlike CHO-K1 cells, CHO-GS cells did not accumulate significant amounts of extracellular alanine (**Fig. 1**), and fed alanine contributed a much larger percentage of the intracellular alanine pool in this cell line (**Fig. 4C**). In CHO-K1, synthesized alanine was seen to be primarily produced from glucose, especially through the pyruvate pool generated directly and indirectly through lactate re-adsorption (**Fig. 4A**).

The individual contributions of the four key amino acids contributing to central metabolism, including the tricarboxylic acid (TCA) cycle, differed significantly for the two CHO cell lines. For CHO-K1 cells, glutamine was initially a major contributor to the citric acid cycle, yielding ∼30-60% of the pools of measured TCA metabolites (**Fig. 4B**). However, its contribution gradually declined during the first three days of culture prior to glutamine being near-fully consumed in the extracellular medium on day 4 (**Fig. 1B**). Labeling from glutamate was also present, but to a lesser degree than from glutamine. As glutamine was depleted, asparagine became a dominant contributor to the citric acid cycle in CHO-K1, peaking on day 4, then decreasing (**Fig. 4B**). The contribution of aspartate, in contrast, remained relatively constant throughout the culture.

CHO-GS cells diverged in their incorporation of the key amino acids due to the overexpression of the *GS* gene and elimination of glutamine from the cell culture media. Asparagine and aspartate replaced glutamine as the major amino acid contributors to the citric acid cycle in early culture (**Fig. 4C**). The contribution of asparagine declined at later days due to its depletion (**Fig. 1C**). Extracellular aspartate was not depleted as rapidly, and the intracellular contribution of aspartate to most TCA cycle intermediates increased to a maximum by day 4 before declining in the final days (**Fig. 4C**). The fate of glutamate also differed in CHO-GS, which exhibited measurable but limited incorporation into TCA cycle intermediates like citrate, malate and fumarate (**Fig. 4C**) especially at earlier days in the culture.

Isotopic labeling data suggests that glycine and serine were rapidly interconverted intracellularly (**Fig. 3**). For both CHO cell lines, extracellular serine was the principal source of both serine and glycine in the cell pellet. Its contribution to both intracellular species began at 80-90% on day 1, gradually decreasing to ∼60% by day 6 (**Fig. 4B, C**). The contribution of extracellular glycine remained relatively constant at about 10-20%, with glucose making up the balance. Serine-glycine interconversion, in addition to methionine labeling from serine and glucose (**Fig. 2**) points to active one-carbon metabolism as will be discussed later.

### 2.3. Mass isotopomer distributions reveal fast-changing exchange fluxes that contribute to rapid turnover of extracellular metabolite pools

The extracellular enrichment of the non-essential amino acids, alanine, glycine, glutamate, serine, and aspartate decreased significantly during amino acid tracer experiments for both cell lines (**Fig. 5**). This decline indicates that these amino acids were actively produced intracellularly by CHO cells from other (unlabeled) sources and then exchanged with the extracellular metabolite pools. To better visualize and quantify the extent of this extracellular metabolite pool turnover, we compared the intracellular and extracellular mass isotopomer distributions of these amino acids over the culture (**Fig. 5A, B**). A rapid decline in extracellular ^13^C-labeling was observed for alanine, glycine, and glutamate, which lost most of their labeling within a few days. For example, by day 2, extracellular alanine concentration increased by 2-fold in CHO-K1 (**Fig. 1B**), but the percentage of extracellular alanine labeling decreased by more than 5-fold over that same interval (**Fig. 5A**). Extracellular serine also shed labeling for both CHO cell lines over time, albeit more gradually (**Fig. 5A, B**). Initially, extracellular serine was exclusively M+3 labeled (that is, all three carbons were labeled), reflecting direct uptake from the media. Over time, partially labeled serine (M+1 and M+2) and unlabeled serine (M+0) were detected intracellularly and extracellularly in both CHO cell lines. Glucose and glycine were the principal sources of M+0 serine (**Fig. 4**) with CHO-K1 generating a higher fraction of this M+0 serine by the end of culture than CHO-GS (**Fig. 5**). This notion of unlabeled and partially labeled serine excretion, particularly in CHO-K1, reflects a notable high pool turnover by day 6 (**Fig. 5A, B**), that occurs despite an overall net consumption of this amino acid (**Fig. 1B, C**).

**Figure 5.**
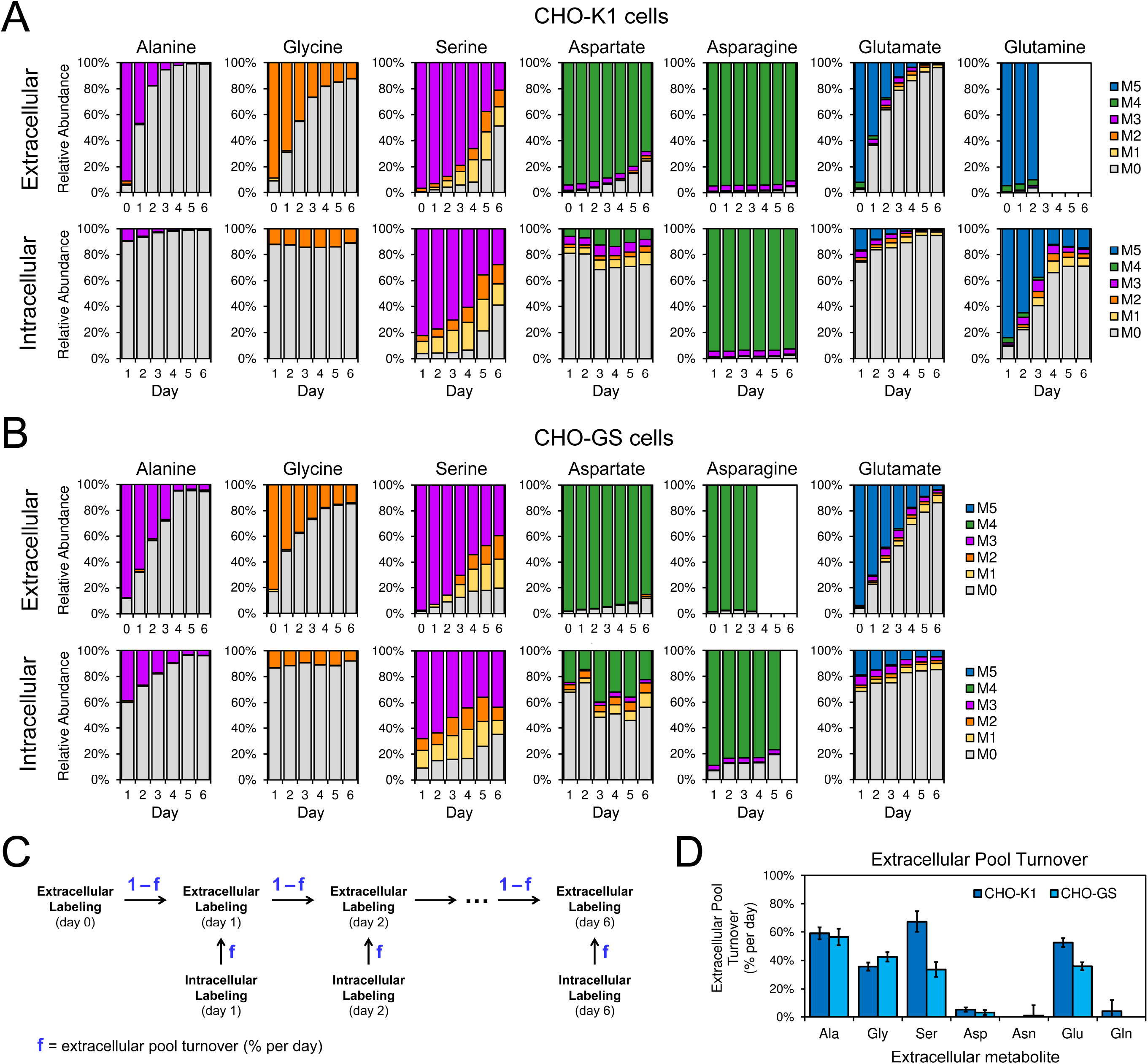
Analysis of extracellular and intracellular isotopic labeling over time identifies turnover of extracellular amino acid pools. Mass isotopomer distributions of extracellular and intracellular amino acids were measured daily for (A) CHO-K1 cells, and (B) CHO-GS cells. Appearance of unlabeled (M0) and partially labeled mass isotopomers in extracellular metabolites indicates that these amino acids were produced from other unlabeled carbon sources. (C) Schematic representation of a mathematical model used to describe turnover of extracellular metabolite pools. The model contains one fitted parameter, f = extracellular pool turnover (% per day). (D) The mass isotopomer data shown in panels A and B were used to estimate extracellular pool turnover rates of amino acids for both CHO cell lines.

We also observed a decrease in the labeling of extracellular aspartate, although to a much lesser degree (**Fig. 5A, B**). By day 6 of the culture, 70% of aspartate was still fully labeled (M+4) in CHO-K1 cell culture, and about 85% fully labeled for CHO-GS cells. At the same time, intracellular aspartate was less than 10% and 25% M+4 for CHO-K1 and CHO-GS cells, respectively. This suggests that the turnover of extracellular aspartate was much slower compared to the turnover of alanine, glycine, serine, and glutamate as will be discussed below. Lastly, we did not observe significant decreases in extracellular labeling for asparagine or glutamine.

Intracellular aspartate showed great isotopomer diversity, with a late-appearing M+1 aspartate in CHO-GS potentially requiring two turns of the TCA cycle. In contrast, M+4 asparagine dominated throughout the culture. The MID for both glutamine and glutamate revealed a progressive shift in the labeled isotopologues over the culture duration, where intracellular M+5 glutamine gradually gave way to M+3, M+2, and finally M+1 by the final days, with a similar pattern mirrored in both intra- and extracellular glutamate across both cell lines (F**ig. 5A, B**). This gradual shift in labeling is consistent with these amino acids cycling through the TCA cycle multiple times and being regenerated with fewer labeled carbons after each successive turn.

To further examine the turnover rate of these extracellular amino acid pools, we implemented a quantitative model with one fitted parameter (*f*) (**Fig. 5C**). In this simple model, extracellular labeling on each day was described as a mixture of intracellular labeling on that day (with contribution *f*) and extracellular labeling carried over from the previous day (with contribution 1-*f*). The fitted parameter in this model (*f*) represents the extracellular pool turnover, expressed as a percentage of the extracellular pool replaced each day. For each tracer experiment, we fitted the measured extracellular labeling to this model using least-squares regression to estimate *f*-values for all amino acids (**Fig. 5D**). The two CHO cell lines exhibited similar turnover rates for most amino acids. The highest turnovers were observed for alanine (∼58% turnover per day), glycine (∼36% per day), glutamate (40-50% per day), and serine (60% per day for CHO-K1 cells, and 30% per day for CHO-GS cells). As expected, the estimated turnover rate for aspartate was low (about 3 to 4% per day), and negligible turnover was estimated for asparagine and glutamine.

The presence of these exchange fluxes has important implications for quantitative stable-isotope analysis methods, particularly ^13^C-metabolic flux analysis (^13^C-MFA). One of the core assumptions of ^13^C-MFA is that the labeling of the tracer remains constant over time. Our data show that this will not be valid for several amino acid tracers. In order apply ^13^C-MFA in the future, more advanced isotopic non-stationary methods will need to be developed and implemented.

### 2.4. Tracing CHO cell metabolic differences with [U-^13^C]glutamine and [U-^13^C]glutamate tracers

Different tracers provide unique windows into cell metabolism; [U-^13^C]glutamine enables valuable insights into the fluxes of the citric acid cycle, glutaminolysis pathway, reductive carboxylation pathway, and pyruvate metabolism (**Fig. 6A**).^24,20,13^ Briefly, if glutamine is metabolized via the glutaminolysis pathway (green arrows in **Fig. 6A**), we expect to observe M+5 labeled glutamate; M+4 labeled succinate, fumarate, malate and oxaloacetate; and M+3 labeled pyruvate and lactate. This M+3 labeled pyruvate can subsequently enter the citric acid cycle via pyruvate carboxylase (PC) to produce M+3 labeled oxaloacetate, malate, and fumarate. If glutamine is instead metabolized via the reductive carboxylation pathway (purple arrows in **Fig. 6A**), we expect to observe M+5 labeled glutamate and citrate, as well as M+3 labeled oxaloacetate, malate and fumarate. Finally, if glutamine is oxidized in the citric acid cycle, M+1 and M+2 labeled glutamate, succinate, fumarate, malate, oxaloacetate, and citrate form, as labeled carbon atoms are lost as ^13^CO_2_ in each subsequent turn of the citric acid cycle.

**Figure 6.**
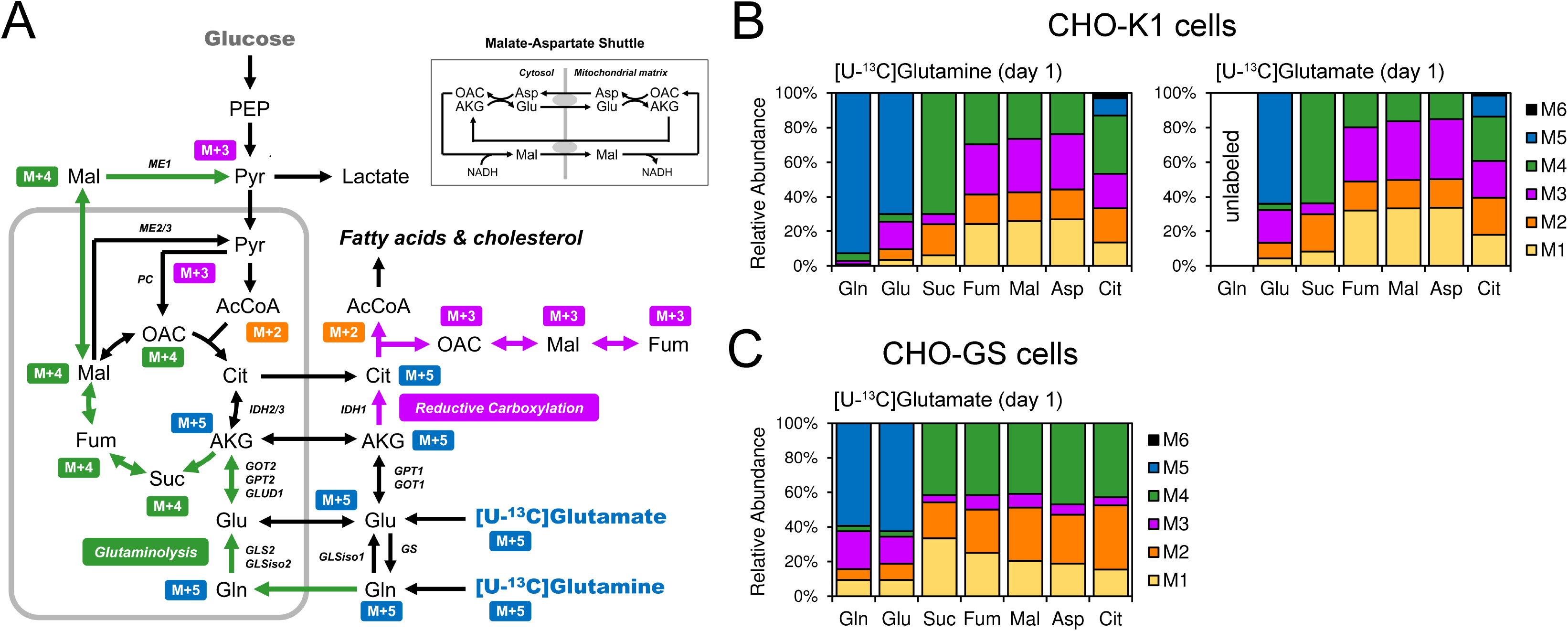
Tracing CHO cell metabolism with [U-^13^C]glutamine and [U-^13^C]glutamate tracers. (A) Schematic of characteristic isotopic labeling patters that can be observed depending on which metabolic pathways are engaged: reductive carboxylation pathway (purple arrows); glutaminolysis pathway (green arrows). (B) Mass isotopomer distributions of intracellular metabolites on day 1 in CHO-K1 cells from [U-^13^C]glutamine and [U-^13^C]glutamate tracer experiments. (C) Mass isotopomer distributions of intracellular metabolites on day 1 in CHO-GS cells from [U-^13^C]glutamate tracer experiment. Measured mass isotopomer distributions were corrected for natural isotope abundances. Only labeled mass isotopomers are shown. All measured labeling data are provided in supplemental materials.

We quantified the intracellular metabolite labeling patterns for CHO-K1 and CHO-GS cells, respectively, during the exponential growth phase (day 1) from tracer experiments with [U-^13^C]glutamine (CHO-K1 cells only) and [U-^13^C]glutamate (both CHO cell lines) (**Fig. 6B, C**).

The differences between the labeling patterns for the two CHO cell lines were significant, demonstrating distinct metabolic physiologies across the cell lines. In CHO-K1, the labeling patterns from [U-^13^C]glutamine and [U-^13^C]glutamate tracer experiments were very similar (**Fig. 6B**); which is not surprising given that the first step in glutamine metabolism is deamination to glutamate. For the tracer experiment with [U-^13^C]glutamate in CHO-K1, we noted that intracellular glutamine was fully unlabeled, indicating that all intracellular glutamine originated from the extracellular (unlabeled) glutamine pool rather than glutamine produced intracellularly. Interestingly, we further observed the presence of M+5 labeled citrate derived from both labeled substrates in CHO-K1, indicating redirection of some glutamine and glutamate via the reductive carboxylation pathway. Consistent with this, we observed high abundances of M+3 labeled aspartate (as a surrogate of oxaloacetate), malate, fumarate, and citrate. For citrate, we also observed that the abundance of M+3 was notably higher than M+5 abundance, which suggests that there was anaplerotic flux from pyruvate to oxaloacetate via pyruvate carboxylase following glutaminolysis and TCA cycle activity. Lastly, a small fraction of M+6 citrate was detected perhaps through a circuitous route likely from the condensation of M+4 oxaloacetate with M+2 acetyl-CoA.

In contrast to CHO-K1, CHO-GS exhibited a very different labeling pattern for glutamate and from fed [U-^13^C]glutamate, consistent with overexpressed amidation reactions via *GS* (**Fig. 6C**). Indeed, we further observed that intracellular glutamine was predominantly M+5 labeled, indicating that extracellular glutamate was the principal source of intracellular glutamine, as compared to lesser amounts of glutamate produced intracellularly in the citric acid cycle. Importantly, no M+5 labeled citrate was detected in this cell line, with very low abundances of M+3 labeled aspartate, malate, fumarate, and citrate (**Fig. 6C**). This distribution indicates that in CHO-GS cells, reductive carboxylation was inactive and there was very limited anaplerotic flux from pyruvate to oxaloacetate. Taken together, our results demonstrate that the CHO-GS cells favor redirection of glutamate flux away from reductive carboxylation and anaplerosis (as observed in CHO-K1 cells) and instead towards glutamine synthesis.

### 2.5. [U-^13^C]Aspartate tracers yield metabolic insights across extended days for batch CHO cell cultures

Glutamine, glutamate, and asparagine (for CHO-GS cells) tracers were limited in following metabolism at later times in batch culture. Glutamine and asparagine were fully consumed by CHO-K1 cells and CHO-GS cells, respectively, within a few days (**Fig. 1**), meaning they could only provide flux information during the early exponential growth phase. Extracellular glutamate lost most of its labeling within a few days (**Fig. 5**), also limiting its use in some cases. To address these concerns and evaluate metabolism at later stages of cell culture, we tracked [U-^13^C]aspartate, which exhibited robust labeling throughout the culture period for both CHO cell lines (**Fig. 3**).

A schematic of relevant metabolic pathways that can impact the observed isotopic labeling patterns in citric acid cycle metabolites from [U-^13^C]aspartate is displayed in **Fig. 7**. Fully ^13^C-labeled aspartate (M+4) can enter the citric acid cycle through one of two metabolic pathways: 1) through the conversion of aspartate to oxaloacetate, as part of the malate-aspartate shuttle that transfers reducing equivalents from cytosolic NADH into the mitochondrial matrix; or 2) through the conversion of aspartate to fumarate, as part of the urea cycle and *de novo* nucleotide biosynthesis pathway. Oxaloacetate, malate and fumarate can be interconverted through the actions of malate dehydrogenase (that interconverts malate and oxaloacetate) and fumarase (that interconverts malate and fumarate); both reactions are known to be reversible. When fully labeled oxaloacetate (M+4) is catabolized in the citric acid cycle, it loses ^13^C-atoms in the decarboxylation reactions (catalyzed by isocitrate dehydrogenase and a-ketoglutarate dehydrogenase), yielding NADH and CO_2_. In the first cycle, this results in M+3 a-ketoglutarate and M+2 succinate; followed by progressively reduced labeled and eventually unlabeled a-ketoglutarate and succinate with an increasing number of TCA cycle rotations (**Fig. 7A**).

**Figure 7.**
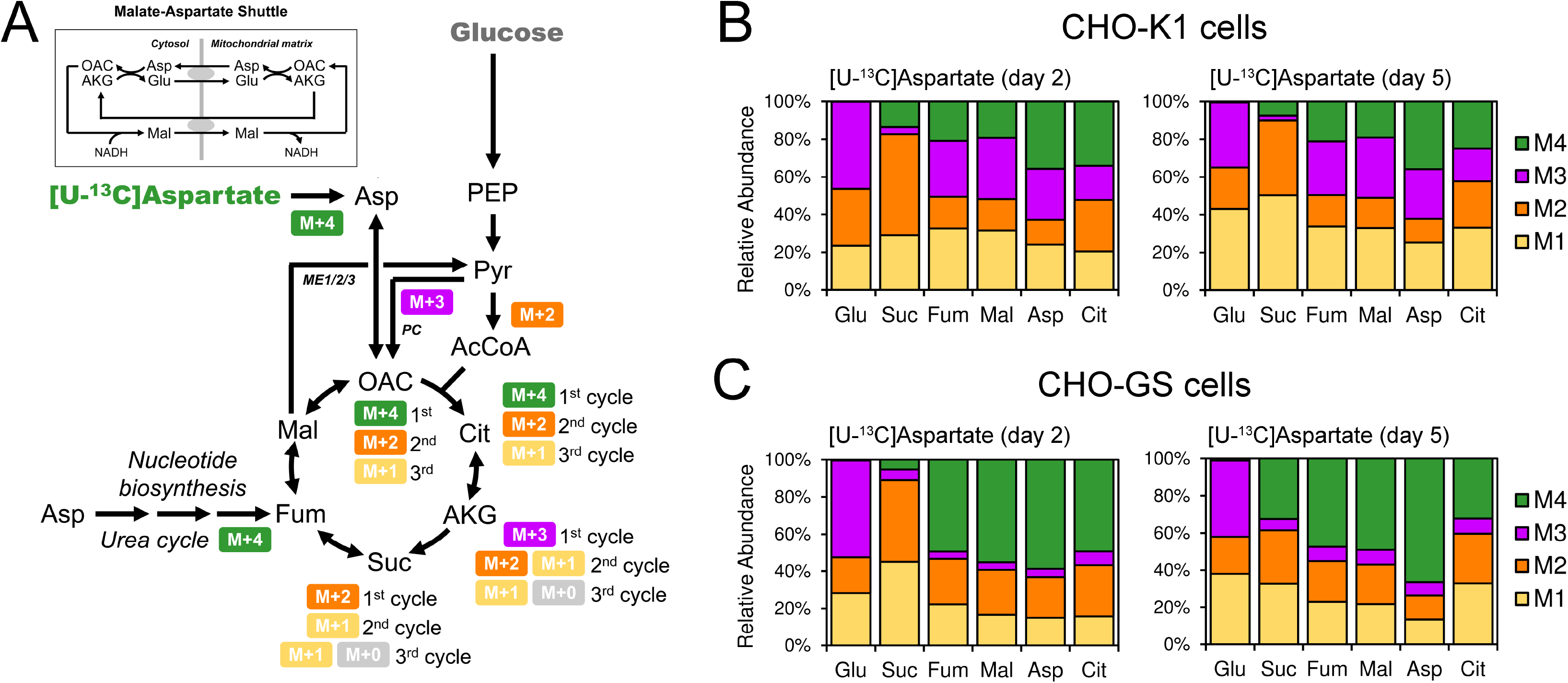
Tracing CHO cell metabolism with [U-^13^C]asparate tracer. (A) Schematic of characteristic isotopic labeling patters that can be observed from metabolism of aspartate in central carbon metabolic pathways. (B) Mass isotopomer distributions of intracellular metabolites on days 2 and 5 in CHO-K1 cells from [U-^13^C]asparate tracer experiment. (C) Mass isotopomer distributions of intracellular metabolites on days 2 and 5 in CHO-GS cells from [U-^13^C]asparate tracer experiment. Measured mass isotopomer distributions were corrected for natural isotope abundances. Only labeled mass isotopomers are shown. All measured labeling data are provided in supplemental materials.

Furthermore, M+3 labeled malate, aspartate and fumarate can also be formed from pyruvate anaplerosis, that is, via the conversion of M+4 malate to M+3 pyruvate (by malic enzyme), followed by the conversion of M+3 pyruvate to M+3 oxaloacetate (by pyruvate carboxylase) (**Fig 7A**). As such, labeling patterns in citric acid cycle metabolites provide critical information regarding the citric acid cycle activity, extent of anaplerosis via pyruvate carboxylase, and total influx of labeled aspartate into the citric acid cycle.

Shown in **Fig. 7B** are the measured labeling patterns for citric acid cycle metabolites and aspartate (which reflects the labeling of oxaloacetate) for [U-^13^C]aspartate tracer experiments with CHO-K1 and CHO-GS cells, respectively. The labeling patterns once again reveal significant metabolic differences between the two CHO cell lines. For example, we observed high abundances of M+3 labeled fumarate, malate, aspartate, and citrate for CHO-K1 cells. This labeling pattern further confirms that anaplerosis from pyruvate is active (**Fig. 7A**), consistent with the results obtained using [U-^13^C]glutamine and [U-^13^C]glutamate tracers (see **Section 2.4**). In contrast, for CHO-GS cells, the M+3 abundances of the same metabolites were low, indicating that anaplerosis from pyruvate was insignificant in CHO-GS cells, again consistent with the result obtained using [U-^13^C]glutamate tracer (see **Section 2.4**). Importantly, the pattern persisted through day 5, which demonstrates that anapleropsis is an inherent physiological characteristic of the CHO-K1 cell line and is not dependent on extracellular glutamine levels. This finding also demonstrates the power of ^13^C-aspartate labeling as a valuable tool to characterize cellular physiology during the later batch phases once glutamine has been depleted and lactate consumption has begun. This anaplerotic activity was further manifested in elevated M+1 abundances for CHO-K1 cells as the oxaloacetate derived from pyruvate traverses the TCA cycle. Alternatively, the CHO-GS cells exhibit elevated M+4 labeling for fumarate, malate, and aspartate, confirming the amplified role of aspartate, as compared to glutamate, feeding into the TCA cycle (either through aspartate-malate shunt or the urea cycle). Indeed, the M+4 labeling of aspartate dominated to an even greater extent on day 5 in CHO-GS because of the increasing importance of aspartate following exhaustion of asparagine.

In contrast, succinate labeling at day 2 for both cell lines was dominated by M+1 and M+2 forms, showing that the flux from M+4 aspartate did not flow reversibly through TCA at fumarate to a large extent. Succinate arose from the forward cycling of the labeled carbons from aspartate through one or multiple cycles of the TCA cycle pathway. The reversible reaction yielding succinate from fumarate was not completely inactive; however, as the M+4 enrichment of succinate reached 30% of all labeled succinate isotopomers on day 5 for CHO-GS cells.

Taken together, our results demonstrate that [U-^13^C]aspartate can provide informative data for differential analysis of metabolism and intracellular dynamics in CHO cells throughout a cell culture. Given that aspartate is not fully consumed by CHO cells during a typical batch cell culture, as opposed to glutamine or asparagine (**Fig. 1**), and the high contribution of aspartate to core metabolism over multiple days (**Fig. 4**), [U-^13^C]aspartate represents an attractive tracer for future metabolic studies in CHO and other mammalian cell cultures.

### 2.6. Fatty acid and cholesterol metabolism in CHO-K1 and CHO-GS cells

Media used for culturing CHO cells typically does not contain non-essential fatty acids or cholesterol ^25^. CHO cells must therefore synthesize these key components of cell membranes *de novo* from glucose or amino acids. Cytosolic acetyl-CoA is the precursor for both fatty acid and cholesterol biosynthesis; as discussed previously, glutamine and glutamate can produce acetyl-CoA via one of two pathways: the reductive carboxylation pathway, or the glutaminolysis pathway (**Fig. 6A**). Asparagine, aspartate, and alanine can also contribute to fatty acid and cholesterol biosynthesis, in part through conversion of citrate into acetyl-CoA from citrate lyase (**Fig. 6A**).

We observed the highest labeling incorporation into fatty acids and cholesterol from [U-^13^C]glucose for both CHO cell lines, although appreciable labeling was also observed in tracer experiments with ^13^C-labeled glutamine (in CHO-K1), with limited inputs from glutamate, asparagine (in CHO-GS), aspartate, and alanine (**Fig. 3**). To quantify the relative contributions of the different carbon sources to fatty acid and cholesterol biosynthesis, we used the isotopomer spectral analysis (ISA) approach to analyze the mass isotopomer distributions measured by GC-MS ^26^. Briefly, ISA is a data analysis technique that estimates two parameters, known as the D and G values. The D value represents the relative contribution of a tracer to lipogenic acetyl-CoA, while the G value represents the fraction of newly synthesized fatty acids (or cholesterol) in the sample. ISA must be used for data analysis to account for the pre-existing (unlabeled) fatty acids and cholesterol present in the samples (see Materials and Methods).

The measured mass isotopomer distributions of palmitate (C16:0) in CHO cell biomass on days 1, 3, and 5, from [U-^13^C]glucose tracer experiments were quantified for both CHO cell lines (**Fig. 8A, B**). Because [U-^13^C]glucose predominantly produced M+2 labeled acetyl-CoA that was then polymerized to produce palmitate (i.e. from 8 acetyl-CoA units), we observed labeling patterns with characteristic even-numbed mass isotopomers. For CHO-K1 cells, the highest abundance mass isotopomers were: M+8 and M+10 (on day 1); M+10 and M+12 (on day 3); and M+14 and M+16 (on day 5). For CHO-GS cells, the highest abundance mass isotopomers of labeled palmitate were: M+10 and M+12 (on day 1); and M+14 and M+16 (on days 3 and 5). These data also indicate that the relative contribution of glucose to lipogenic acetyl-CoA increased over time. The fraction of unlabeled palmitate (M+0), that is, palmitate from the inoculum, decreased over time for both cell lines as was expected.

**Figure 8.**
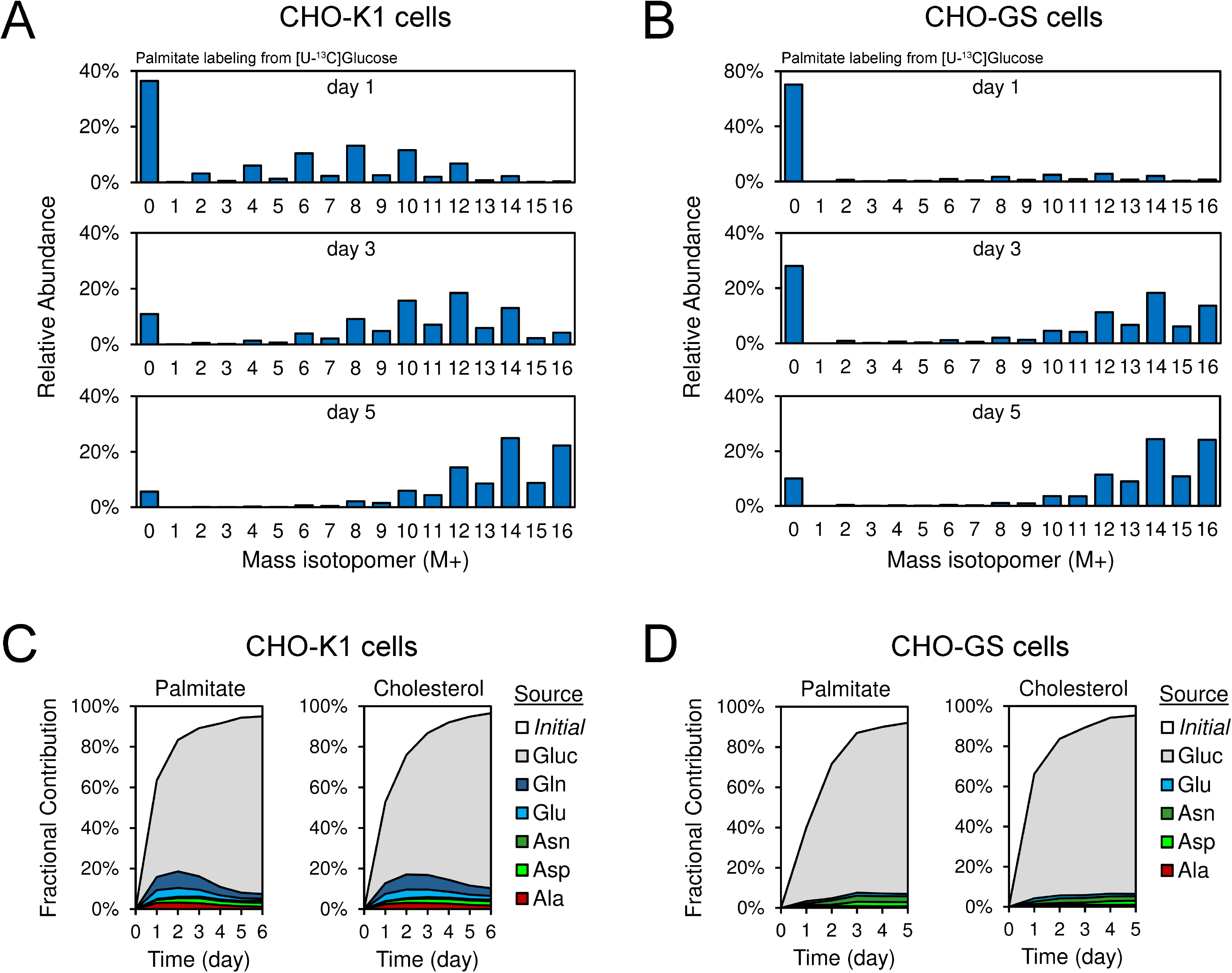
Fatty acid and cholesterol metabolism in CHO-K1 and CHO-GS cells elucidated with ^13^C-tracers. Mass isotopomer distributions of palmitate measured on days 1, 3 and 5 in tracer experiments with [U-^13^C]glucose for (A) CHO-K1 cells, and (B) CHO-GS cells. The measured mass isotopomer distributions were corrected for natural isotope abundances. Measured labeling data on all days are provided in supplemental materials. The isotopic labeling was analyzed using the ISA approach to calculate fractional contributions of different carbon sources to the synthesis of fatty acids and cholesterol for (C) CHO-K1 cells, and (D) CHO-GS cells on different days of culture.

Using the results from ISA, we calculated the relative contributions of the different sources to palmitate and cholesterol over time for both CHO cell lines (**Fig. 8C, D**). For CHO-K1 cells, we observed that during the early exponential growth phase (days 0-2), glucose contributed about 60% to lipogenic acetyl-CoA, with the remaining 20% coming from a combination of different amino acids, including glutamine and glutamate primarily through reductive carboxylation (**Fig 6B).** Additional contributions were likely from glutaminolysis, as well as from aspartate, asparagine, and alanine. The contribution of alanine for lipid biosynthesis is consistent with its excess availability (**Fig. 1**) as a side effect of the ammonium generation from glutamine deamination in CHO-K1. After day 3, the contribution of glucose increased with lower contributions to acetyl-CoA originating from the amino acids, coinciding with the exhaustion of glutamine. In contrast, for CHO-GS cells, the contribution of amino acids to lipogenic acetyl-CoA was lower for most of the culture period, with at most ∼7% coming from amino acids (**Fig. 8D)** primarily through glutaminolysis and anaplerosis rather than reductive carboxylation. This indirect route to acetyl-CoA resulted in lesser contributions of glutamate to palmitate and cholesterol as compared to CHO-K1, with asparagine a contributor to palmitate and cholesterol biosynthesis in CHO-GS cells.

### 2.7. Elucidating one-carbon metabolism in CHO cells

Finally, we analyzed the labeling patterns of glycine, serine, and methionine to gain insights into one-carbon metabolism in the different CHO cells (**Fig. 9**). Previous studies in CHO cells have lacked experimental data to determine one-carbon flows across the relevant metabolic pathways, including the folate cycle, methionine cycle, and trans-sulfuration pathway. Together, these reaction networks are crucial for a variety of cellular processes such as dNTP synthesis, purine synthesis, GSH generation and DNA-histone methylation (**Fig. 9A**). While transcriptomic studies have been conducted for determining the activity of several genes in the trans-sulfuration branch of the one-carbon flow ^27^, the dynamics of one-carbon metabolism in CHO cells remain largely unexplored.

**Figure 9.**
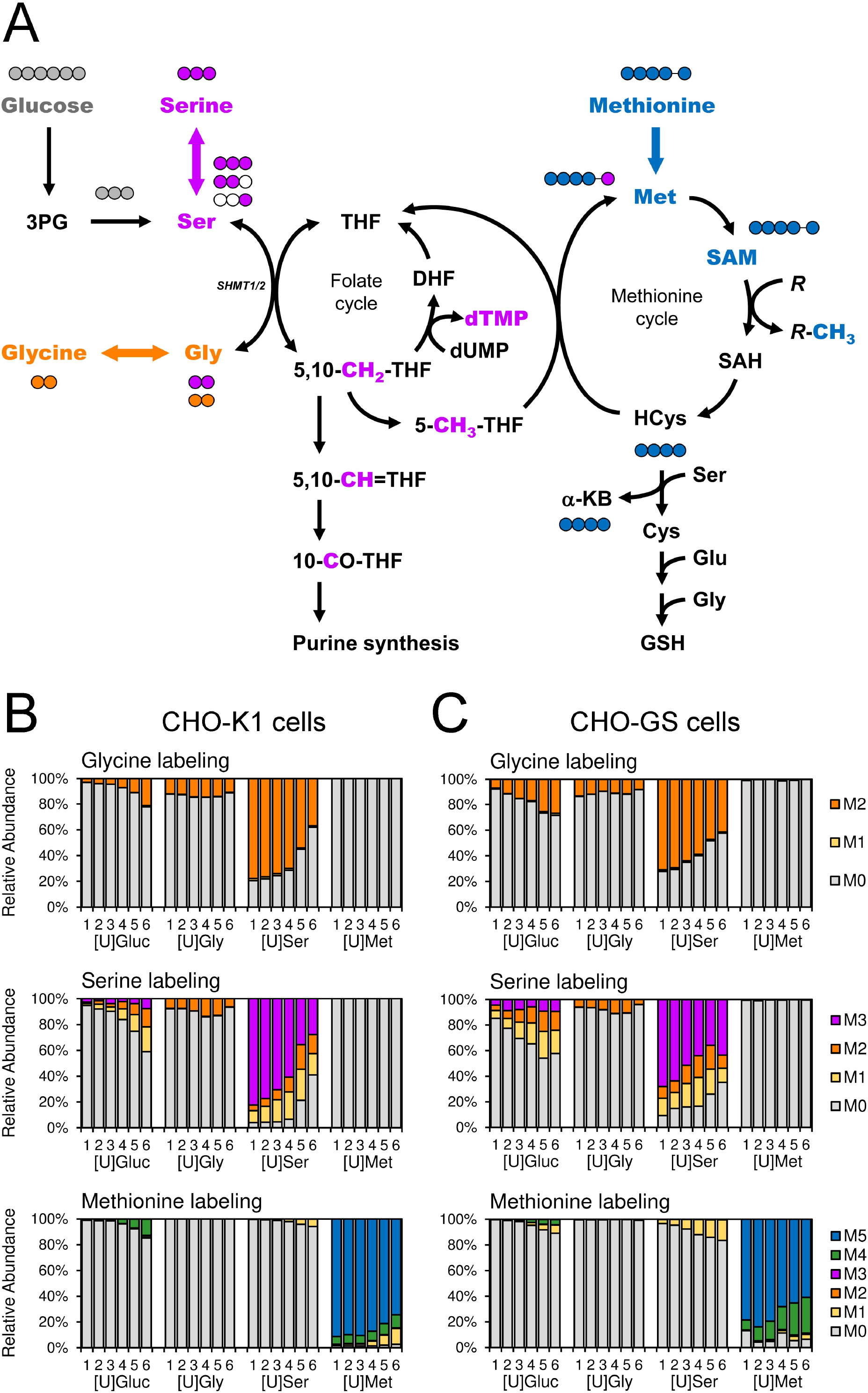
One-carbon metabolism in CHO cells elucidated with ^13^C-tracers. (A) Schematic of metabolic pathways involved the production and consumption of one-carbon units derived from glucose, serine, glycine, and methionine. Shown are the measured mass isotopomer distributions of intracellular glycine, serine and methionine in tracers experiments with [U-^13^C]glucose, [U-^13^C]glycine, [U-^13^C]serine, and [U-^13^C]methionine for (B) CHO-K1 cells, and (C) CHO-GS cells. Mass isotopomer distributions were corrected for natural isotope abundances. All measured labeling data are provided in supplemental materials.

Mass isotopomer distributions of intracellular glycine, serine, and methionine were generated from tracer experiments with ^13^C-labeled glucose, glycine, serine, and methionine (**Fig. 9B, C**). As previously noted, glycine and serine were derived primarily from extracellular serine in both cell lines (see Sections 2.2 and 2.3). Serine hydroxymethyltransferase (SHMT) is the crucial enzyme in the folate-mediated one-carbon metabolism that catalyzes the reversible conversion of serine to glycine, generating 5,10-methylenetetrahydrofolate (5,10-CH_2_-THF), which is essential for DNA synthesis, repair, and cell proliferation. The reversibility of the SHMT reaction is evident from the labeling patterns we observed for intracellular serine from tracer experiments with [U-^13^C]serine and [U-^13^C]glucose. Specifically, we observed high abundances of M+1 and M+2 labeled serine in both CHO cell lines. These mass isotopomers are formed when M+3 labeled serine is first converted to M+2 labeled glycine and M+1 labeled 5,10-CH_2_-THF by SHMT, followed by the reverse reaction and back again (i.e. glycine + 5,10-CH_2_-THF to serine + THF) (**Fig. 9A**). Evidence of this pathway was also indicated by the M+2 glycine pool formed from [U-^13^C]serine and [U-^13^C]glucose. The fractional decline of the M+2 glycine pool derived from serine over days is likely due in part to increasing contributions to the glycine pool from unlabeled glucose (**Fig. 9B, C**). When comparing the labeling patterns for the two CHO cell lines, we also observed a lower contribution of glucose to serine in CHO-K1 cells compared to CHO-GS cells on days 0-4.

The labeling patterns of intracellular methionine provided additional insights into the activity of the methionine cycle and one-carbon flow from the folate cycle to the methionine cycle (**Fig. 9A**). In this cycle, methionine is first activated into S-adenosylmethionine (SAM), which then donates a methyl group to various biological targets (e.g. DNA, RNA, proteins, lipids) and becomes S-adenosylhomocysteine (SAH). SAH is next converted to homocysteine; and finally, to complete the cycle, homocysteine is converted back into methionine by acquiring a methyl group from the folate cycle. When CHO-GS cells were cultured with [U-^13^C]methionine, we initially observed predominantly M+5 labeled methionine, which gradually declined from about 80% to approximately 60% over time. During the same time, the abundance of M+4 methionine gradually increased from 8% to 28%, indicating that the M+5 labeled methionine donated its ^13^C-labeled methyl group over time and gained an unlabeled methyl group from the folate cycle. This was further supported by the corresponding appearance of M+1 labeled methionine in tracer experiments with [U-^13^C]serine (**Fig. 9C**), given that serine was the major contributor of one-carbon units to the methionine cycle. Interestingly, for CHO-K1 cells, we observed much lower methionine cycle activity in terms of M+4 labeling over the duration of the batch culture. Furthermore, methionine showed almost no M+1 labeling over the first 3 days when using [U-^13^C]serine tracer Only in the last three days of the culture (days 4 to 6) did we observe evidence of active low level serine contributions to the adjacent methionine cycle and serine’s M+1 contribution to methionine never increased above 6% in CHO-K1 while representing 16% in the CHO-GS (**Fig. 9B**), pointing to a reduced rate of methionine cycling in the CHO-K1 cell line.

## 3. Discussion

### 3.1. Overview of the ¹³C-based metabolic framework

Our framework integrated tracer experiments with GC-MS based isotopomer analysis for time-resolved ^13^C-enrichment, enabling mapping of carbon flows from substrate contributions to specific metabolic intermediates and biomass components. This approach provides a dynamic, systems-level view of central carbon metabolism that extends beyond static measurements of metabolite concentrations previously available. Moreover, combining ^13^C-enrichment analysis with mass isotopomer distribution analysis for each metabolite across cell lines and timepoints yields insights concerning the activity of pathways emerging from central carbon metabolism and how they differ across the most important production CHO cell lines. We also employed isotopomer spectral analysis (ISA) of GC-MS-derived mass isotopomer distributions to quantitatively partition the contributions of glucose and multiple amino acids to lipogenic acetyl-CoA, providing a more comprehensive picture of fatty acid and cholesterol biosynthesis than has been previously reported. Also, to our knowledge, this study provides the first ever analysis of one-carbon metabolism utilizing select ^13^C amino acid and glucose tracers in CHO, indicating differences in folate and methionine metabolism across the two CHO cell platforms.

Distinct differences in metabolic physiologies emerged when comparing ^13^C-glucose and amino acid labeling patterns between these cell lines, due to the selection strategies used to generate them: CHO-K1 cells were previously selected using methotrexate to driving DHFR gene amplification, while CHO-GS cells were selected using the glutamine synthetase gene ^8^.

The specific cell line selection strategy used imparts significant changes in cellular physiology spanning key metabolic and energy pathways. Broadly speaking, we show that four key non-essential amino acids — glutamate, glutamine, aspartate and asparagine (with alanine to a lesser extent) — along with glucose, were the primary drivers for central carbon metabolic activity while essential amino acids were largely isolated from central carbon metabolism. The synthesis of glycolytic and TCA cycle intermediates, and even cellular fatty acids and cholesterol from these substrates differed across cells lines, with ^13^C-tracing demonstrating how substrate contributions to crucial metabolites varied over time.

### 3.2. TCA cycle anaplerosis and overflow metabolism

Individual ^13^C amino acid labeling revealed fundamentally different metabolic patterns between CHO-K1 and CHO-GS especially for the TCA cycle. In CHO-K1 cells, glutamine was the largest amino acid contributor to TCA cycle activity through glutaminolysis-driven generation of succinate, fumarate, and malate, and further contributed to amino acid pools (glutamate, aspartate), glycolytic products (pyruvate, lactate), and lipids (palmitate, cholesterol). Most importantly, CHO-K1 cells engaged in extensive reductive carboxylation and active pyruvate anaplerosis via pyruvate carboxylase, neither of which were evident in CHO-GS cells. This activity is consistent with the Warburg Q effect or glutamine respirofermentation, in which rapidly proliferating cancer and immune cells ^28^ utilize high amounts of glutamine and excrete metabolic by-products such as lactate even when oxygen is available. This glutamine overflow metabolism occurs when metabolic demands exceed the mitochondrial capacity to oxidize glucose and glutamine efficiently and in cancers in which mitochondrial TCA functions or electron transport chain is insufficient or dysfunctional ^29,30^. Rather than maximizing ATP yield per molecule of substrate, cells prioritize biomass accumulation by providing precursors for proteins, lipids, and nucleotides and maintaining redox balance ^31,32^. This metabolic phenotype is consistent with our findings of glutamine contributing to secreted alanine, citrate, and lactate in addition to contributing to intracellular pools of other metabolites.

CHO-GS, which lacked supplemental glutamine due to overexpression of glutamine synthase, preferentially redirected glutamate flux toward glutamine synthesis with no evidence of reductive carboxylation or substantial pyruvate carboxylase activity. In the absence of glutamine, CHO-GS relied on asparagine as the primary anaplerotic substrate in early culture ^19,33^, with aspartate filling the role of carbon and nitrogen supplier for the TCA cycle only after asparagine exhaustion. Thes differences in phenotypess between cell lines was not simply a consequence of glutamine availability, since [U-¹³C]aspartate labeling revealed the same metabolic differences across the two cell lines even after glutamine was consumed. This change in metabolic profiles may be due, at least in part, to the overexpression of GS in CHO-GS and the prioritization of glutamate for glutamine synthesis.

### 3.3. De novo biosynthesis: asparagine and alanine

While asparagine was used as a primary initial carbon source in CHO-GS in order to fuel macromolecular, purine, and pyrimidine synthesis, glucose also contributed to its active de novo synthesis later in culture, likely via asparagine synthase (ASNS). In contrast, CHO-K1 maintained near-100% asparagine labeling — consistent with ASNS suppression by excess glutamine and negative regulation by elevated intracellular asparagine concentrations ^19,34^.

Alanine also exhibited a markedly different accumulation pattern between cell lines, with significant extracellular secretion in CHO-K1 versus limited net uptake in CHO-GS over the first 4 days. In CHO-K1, ammonia from glutamine deamidation was likely disposed of via transamination of glutamate and pyruvate to alpha-ketoglutarate and alanine (via ALT), with alanine secreted as a nitrogen sink ^35,36^ — while in CHO-GS, the glutamate pool was instead directed toward glutamine synthesis by GS gene expression, limiting alanine production and forcing reliance on exogenous alanine. Nonetheless, alanine represented another amino acid rapidly exchanged between intracellular and extracellular environments in both hosts.

### 3.4. The Malate-aspartate shuttle and redox balance

Interestingly, fed aspartate contributed a larger fractional share to TCA metabolite pools in CHO-GS than asparagine. Aspartate contributed significantly to intracellular fumarate and malate, particularly in CHO-GS, reflecting its adjacency to these metabolites through oxaloacetate via aspartate aminotransferase (AST) ^37^; this contribution was especially prominent during the mid- to late exponential phase and was channeled through the malate-aspartate shuttle or urea cycle. Utilization of these shuttles in CHO-GS was indicated by the large fractional contribution of aspartate to M+4 isotopologues of fumarate, malate, and aspartate on days 2 and 5. This substantial contribution to intracellular malate and fumarate was unexpected in CHO-GS given aspartate’s relatively slow consumption rate, suggesting that its primary importance lies in shuttling reducing equivalents into the mitochondria rather than as a direct carbon and nitrogen source when asparagine and glutamine are available at sufficient concentrations. Enhanced malate-aspartate shuttle activity, arising from robust intracellular aspartate availability ^38^, can accelerate cytosol-to-mitochondria NADH transfer (**Fig. 6A**), supplying electrons to the ETC linked to ATP generation ^39^. Alternatively, M+4 labeling was considerably less prevalent in CHO-K1, where glutamine consumption was an important NADH source; intracellular aspartate stores in CHO-K1 were additionally supplemented via pyruvate carboxylase activity in this cell line. However, in both cell lines, aspartate showed limited involvement in succinate accumulation, consistent with the large equilibrium constant of succinate dehydrogenase favoring conversion to fumarate over the reverse reaction ^40^. In addition, considerable pools of intracellular aspartate were also obtained from glucose, glutamine (in CHO-K1), glutamate, and asparagine, rather than derived from fed aspartate alone. Enrichment from other sources was higher in CHO-K1 than CHO-GS, likely due to the enhanced glutaminolysis and subsequent asparagine availability. This conserved strategy to supplement direct aspartate uptake with other sources, also observed in cancer cell metabolism ^41^, is likely to be critical for maintaining redox state and precursor pools needed for sustaining nucleotide and protein synthesis.

As a result of its relatively slow consumption rate, aspartate represents a valuable alternative tracer compared to glutamine and asparagine, which are consumed preferentially in CHO cell lines. While [U-^13^C]glutamine and [U-^13^C]glutamate tracers have been widely used for metabolic flux analysis in CHO cells, their utility is inherently limited to the early exponential phase, as glutamine is rapidly depleted and extracellular glutamate loses most of its labeling within a few days. Asparagine, though useful for CHO-GS cells, faces a similar limitation. In contrast, we show here that [U-^13^C]aspartate maintains high intracellular labeling throughout the entire culture duration in both cell lines, making it a uniquely powerful tracer for characterizing metabolism during the late exponential and stationary phase when other tracers are no longer informative. Indeed, the observation of significant M+5 pools of citrate and M+3 pools of multiple TCA cycle intermediates (Fum, Mal, Cit) provide evidence of robust reductive carboxylation and pyruvate carboxylase activity, respectively, in CHO-K1 cells even following the exhaustion of glutamine. In this way, [U-^13^C]aspartate represents as an emerging tool for investigating metabolic fluxes in mammalian lines following glutamine and asparagine exhaustion.

### 3.5. Lipid biosynthesis and carbon source distribution

Applying ISA across multiple tracers revealed the differences in amino acid contributions to lipogenic products — limited at most ∼7% throughout culture in CHO-GS versus a maximum of almost 20% in CHO-K1 — also reflects the broader metabolic rewiring imposed by glutamine availability versus GS amplification. In CHO-K1, early engagement of reductive carboxylation provided a more direct route for glutamine and glutamate to contribute to fatty acids, accounting for higher amino acid contributions to lipid biosynthesis during the early exponential phase; the lack of activity for this pathway in CHO-GS, combined with the lack of glutamine, forced near-exclusive reliance on glucose for lipid synthesis from the outset. Consequently, glucose contributions to palmitate were higher in CHO-GS at day 1 due to fewer alternative precursor sources, but by day 5 glucose dominated palmitate labeling in both cell lines following glutamine exhaustion in CHO-K1. These differences underscore how glutamine availability and differential cell engineering approaches fundamentally shape the broader biosynthetic landscape of both cell lines.

### 3.6. One-carbon metabolism: the folate and methionine cycles

Serine and glycine support the folate cycle as major carbon donors in dNTP generation. A diverse labeling pattern was observed for serine: the M+3 isotopomer reflected direct uptake; the M+2 isotopologue arose from serine donating its beta-carbon to the folate cycle to generate M+2 glycine, which was converted back to M+2 serine using an unlabeled one-carbon units; and M+1 serine resulted from labeled serine-derived carbon recombining with unlabeled glycine, or after two cycles through the folate cycle. Glucose, and to a lesser degree glycine, were the principal sources of unlabeled serine, and some labeled intracellular serine was secreted back into the culture broth, yielding a broad extracellular labeling pattern. Due to methotrexate selection, CHO-K1 cells contained elevated DHFR gene copies and enzyme activity, predisposing them to increased recycling through the inner folate cycle; this gene amplification likely leads to enhanced serine turnover in CHO-K1, where approximately 65% of the extracellular pool of serine is turned over per day as compared to less than 40% in CHO-GS.

In CHO-GS cells, active methionine cycling is evident, with the progressive loss of the labeled methyl groups from M+5 methionine and its replacement with an unlabeled one-carbon units from the folate cycle, corroborated by the progressive appearance of M+1 methionine in serine tracer experiments. In CHO-K1 cells, methionine cycle activity was limited during the first four days, only becoming evident in the serine tracing on the final days of culture. This reduced cycling in the outer cycle in CHO-K1 is consistent with its methotrexate selection, which through DHFR gene amplification would confer a greater intrinsic capacity to generate tetrahydrofolate through the inner folate cycle, thereby reducing the cell’s dependence on active recycling through the methionine cycle. Together these findings indicate that the two cell lines have very different one-carbon metabolic pathway fluxes, with implications for their respective capacities for methylation, nucleotide synthesis, and redox balance. Thus, this isotope tracing framework enables for the first time a dynamic characterization of one-carbon metabolism in CHO cells, encompassing the folate cycle, methionine cycle, and their interconnections with serine, glycine, and methionine metabolism, pathways that have remained largely uncharacterized in CHO despite their critical roles in dNTP synthesis, purine biosynthesis, glutathione generation, and epigenetic regulation.

## 4. Conclusions

In this study, we established a comprehensive and systematic ^13^C-tracing framework capable of mapping the fate of individual nutrients, particularly amino acids and glucose, from extracellular uptake to their incorporation into intracellular metabolites, biomass, and secreted metabolites for mammalian cell lines. By replacing each substrate with its fully labeled counterpart across parallel experiments, we achieved high-resolution interrogation of metabolic pathway utilization in different commercially important CHO cell lines and their corresponding media. Through this approach, we elucidated cell line-specific metabolic phenotypes for CHO-K1 and CHO-GS cells. The physiological characterization presented here highlights both conserved features of CHO metabolism such as glucose consumption and lactate switch, as well as differential dependencies on glutamine, glutamate, asparagine, aspartate, and other amino acids. Indeed, specific CHO cell lineages will require formulations optimized based on their metabolic phenotype which can vary greatly due to selection criteria as well as clonal isolation; thus, a one size fits all media formulation does not “fit” for optimal cell line performance and rebalancing of specific amino acids may be required. Furthermore, our enhanced knowledge about which metabolic pathways are utilized in different CHO lineages will further inform the tailored design of more appropriate cell line specific mathematical models for process monitoring and control. Deficiencies and overactivity can also drive directed metabolic engineering targets to upregulate underutilized pathway or suppress elevated ones. Together, these results establish a foundation into how tracer-based analyses can be applied to decipher insights into cell-specific nutrient and metabolic partitioning as well as changes in overall cellular physiology, knowledge critical for the optimization of biomanufacturing processes needed to produce our emerging biologics in the future.

## 5. Materials and methods

### 5.1. Materials

Media and chemicals were purchased from Sigma-Aldrich (St. Louis, MO). A chemically defined CHO cell culture medium was used in this study (Sigma-Aldrich product no. 87093C). In total, 21 formulations of this medium were purchased, where in each formulation a different amino acid or glucose was omitted. The following tracers were purchased from Cambridge Isotope Laboratories: [U-^13^C]cysteine, [U-^13^C]histidine, [U-^13^C]methionine, [U-^13^C]phenylalanine, [U-^13^C]threonine, [U-^13^C]tryptophan, [U-^13^C]valine, [U-^13^C]leucine, [U-^13^C]aspartate, and [U-^13^C]glucose. The following tracers were purchased from Isotec/Sigma-Aldrich (St. Louis, MO): [U-^13^arginine, [U-^13^C]asparagine, [U-^13^C]lysine, [U-^13^C]proline, [U-^13^C]tyrosine, [U-^13^C]serine, [U-^13^C]alanine, [U-^13^C]glycine, [U-^13^C]isoleucine, [U-^13^C]glutamate, [U-^13^C]glutamine. To prepare ^13^C-labeled media, an appropriate amount of ^13^C-amino acid or ^13^C-glucose was added to the medium that was missing the respective compound and the media were then sterilized by filtration.

### 5.2. Cell lines and culture conditions

Two CHO cell lines were used in this study, a CHO-K1 cell line producing IgG provided by the National Institute of Health (NIH), and a GS-CHO (CHOZN23) cell line producing IgG provided by Millipore-Sigma. CHO cells were cultured in suspension culture in 125 mL shake flasks with vented caps (Corning no. 431143) with 30 mL working culture volume, at 125 rpm shaking speed in a humidified incubator at 37 °C and 5% CO_2_. To prevent cell clumping in CHO-K1 cultures, 100 µL of an anti-clumping agent (Gibco no. 0010057AE) was added to the medium. For CHO-K1 cultures, the medium contained 6 mM glutamine. For GS-CHO cultures, no glutamine was present in the medium. For the tracer experiments, cultures were seeded at 0.3×10^6^ cells/mL and medium samples and cell pellets were collected daily for GC-MS analysis. To obtain cell pellets and cell-free spent media, a predetermined amount of cell culture (∼3×10^6^ cells) was harvested and centrifuged at 1000 rpm for 10min to separate the supernatant from cell pellets. Spent media were filtered and stored at −20 °C. Cell pellets were washed once using 4 °C saline solution or DPBS and stored at −20 °C. The wash step is essential to ensure high resolution in the stable-isotope labeling data acquisition, since any residual labeling from the media can result in inaccurate measurement of enrichment in intracellular metabolites.

### 5.3. Analytical methods

Glucose and lactate concentrations were determined using a YSI 2700 biochemistry analyzer (YSI, Yellow Springs, OH). For the CHO-K1 cultures, VCD was quantified using a Moxi Z cell counter. For GS-CHO cultures, VCD was quantified using a hemocytometer. Cell viability was above 95% in all batch cultures, as determined by Trypan blue staining. For quantification of extracellular amino acid concentrations, a ^13^C-labeled internal standard solution, containing [U-^13^C]algal amino acids with known concentrations, was added to medium samples prior to GC-MS analysis^42^.

### 5.4. Extraction of intracellular metabolites, fatty acids and cholesterol

Intracellular metabolites, fatty acids and cholesterol were extracted as described previously^15,43^. Briefly, 0.4 mL of cold methanol (−20 °C) was first added to the cell pellets, and the tubes were vortexed vigorously for 10 sec. Next, 0.3 mL of chloroform was added and the tubes were vortexed again vigorously for 10 sec. Lastly, 0.3 mL of DI water was added and the tubes were vortexed vigorously for 10 sec. The tubes were then centrifuged at 14,000 rpm and 4 °C for 10 min. The resulting phase separation produced an aqueous upper phase containing intracellular metabolites, and an organic lower phase containing fatty acids and cholesterol.

### 5.5. Gas chromatography mass spectrometry (GC-MS)

GC-MS analysis was performed on an Agilent 7890A GC system equipped with a DB-5MS capillary column (30 m, 0.25 mm i.d., 0.25 μm-phase thickness; Agilent J&W Scientific), connected to an Agilent 5977B Mass Spectrometer operating under ionization by electron impact (EI) at 70 eV. Helium flow was maintained at 1 mL/min. The source temperature was maintained at 230 °C, the MS quad temperature at 150 °C, the interface temperature at 280 °C. For GC-MS analysis of MOX-TBDMS derivatized amino acids and intracellular metabolites^44^, the inlet temperature was 280 °C; 1 μL of derivatized sample was injected at a split ratio based on the peak intensities, typically split 4 or 10. The column was started at 80 °C for 2 min, increased to 280 °C at 7 °C/ min, and held for 20 min. For GC-MS analysis of cholesterol and fatty acid methyl esters (FAME)^15^, the inlet temperature was 250 °C; 1 μL of derivatized sample was injected in splitless mode. The column was started at 80°C for 2 min, increased to 280°C at 10°C/min, and held for 12 min. All mass spectra were recorded in single ion monitoring (SIM) mode with 4 ms dwell time on each ion. Mass isotopomer distributions were obtained by integration of ion chromatograms and corrected for natural isotope abundances^45^.

## Acknowledgements

This work was funded and supported by the Advanced Mammalian Biomanufacturing Innovation Center (AMBIC) through the Industry-University Cooperative Research Center Program under U.S. National Science Foundation grant number 1624684.

## Author Contributions

**V.G.D.:** Investigation, Validation, Formal analysis, Data curation, Writing – original draft, Writing – review & editing. **J.E.G.:** Investigation, Formal analysis, Data curation, Validation. **H.M.N.:** Investigation, Formal analysis, Validation, Writing – original draft, Writing – review & editing. **B.O.M.:** Investigation, Formal analysis, Validation. **P.K.:** Writing – original draft, Writing – review & editing. **J.S.:** Writing – original draft, Writing – review & editing. **M.R.A.:** Conceptualization, Writing – original draft, Writing – review & editing, Data curation, Visualization, Supervision. **M.J.B.:** Conceptualization, Funding acquisition, Writing – original draft, Writing – review & editing, Data curation, Project administration, Supervision.

